# Chronic diazepam reveals excessive homeostatic gain in SOD1^G93A^ mouse spinal motoneurons

**DOI:** 10.64898/2026.05.16.725609

**Authors:** Emily Reedich, Yi-Tzai Chen, Rebecca Imhoff-Manuel, Deyu Li, Marin Manuel

## Abstract

Motoneurons are under strong pressure to maintain stable motor output throughout an individual life, through homeostatic regulation of their electrical properties. Dysregulated spinal motoneuron excitability has long been implicated in the pathogenesis of amyotrophic lateral sclerosis (ALS). Recent work in SOD1^G93A^ mice suggests that the homeostatic response of motoneurons becomes dysregulated as cellular processes are disrupted by the disease, causing fluctuations in motoneuron electrical properties. Yet, few studies directly test whether ALS motoneurons respond differently than wild type motoneurons to a common chronic perturbation. Here, we used in vivo electrophysiology to test whether motoneurons from pre-symptomatic SOD1^G93A^ mice modulate excitability differently than wild type motoneurons in response to the same homeostatic perturbation: chronic inhibition exerted by the benzodiazepine diazepam. Using linear mixed-effects statistical models, we assessed whether diazepam treatment differentially modulated passive properties, firing behavior, spike properties, and/or synaptic inputs in SOD1^G93A^ versus wild type motoneurons. We identified a significant genotype × treatment interaction effect selectively for properties related to passive membrane integration and spike initiation, including membrane time constant, peak input resistance, and recruitment current. In contrast, firing gain, spike waveform characteristics, and synaptic inputs were largely unaffected. These findings indicate that sustained inhibitory perturbation selectively triggered overactive intrinsic compensatory mechanisms in SOD1^G93A^ motoneurons rather than inducing widespread changes in firing or synaptic transmission. Together, our results provide direct evidence for over-active homeostatic control of motoneuron excitability and support a view of motoneuron dysfunction in ALS as a problem of altered feedback regulation rather than simply hyper- or hypo-excitability.

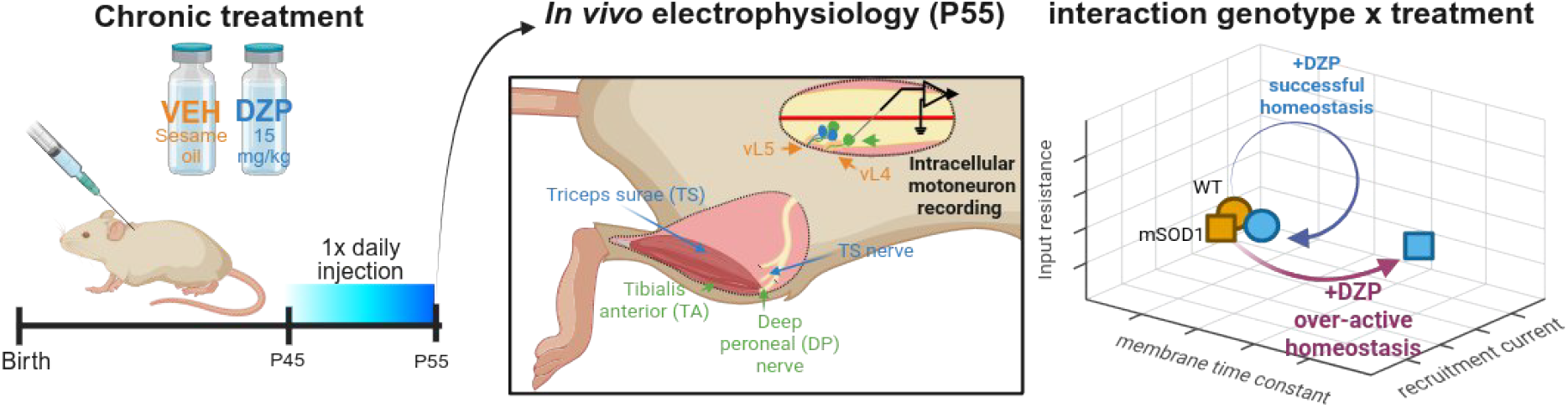

## 1. Introduction

Amyotrophic lateral sclerosis (ALS) is a fatal neurodegenerative disease caused by progressive upper and lower motoneuron degeneration. Altered motoneuron excitability has been implicated in ALS pathogenesis for decades, originally framed within the excitotoxicity hypothesis, linking elevated firing and glutamatergic drive to neuronal death [1], [2]. Recent work suggests a more nuanced pathophysiology, where motoneuron excitability, in a pattern dependent on motoneuron physiological type, oscillates throughout disease progression. In mouse models and human induced pluripotent stem cell-derived motoneurons, early disease stages are often characterized by motoneuron hyperexcitability [3], [4], [5], [6], [7], reflected by reduced rheobase, increased frequency-current (F–I) gain, and enhanced persistent inward currents (PICs). A short-lived period of normal excitability due to compensatory increases in cell size and increased input conductance [8], [9], [10] is followed at later stages by motoneuron hypoexcitability or firing failure [11], [12]. This oscillatory pattern likely reflects dysregulated homeostatic control of motoneuron firing output to maintain motor function during pre-symptomatic stages, in the face of a myriad of disrupted cellular processes.

Motoneurons are equipped with strong intrinsic and synaptic homeostatic mechanisms to stabilize firing output despite large perturbations in aspects like conductance, morphology, and synaptic input [13], [14]. Experimental and computational work has shown that changes in PICs, leak conductance, input resistance, and cell size are often co-regulated in a manner consistent with output preservation rather than runaway excitability [15], [16]. In ALS models, homeostatic mechanisms appear to remain active but operate with excessive gain, producing large compensatory changes in response to modest perturbations [17], [18]. Time-course analyses in the SOD1^G93A^ mouse model of ALS (mSOD1 hereafter) reveal oscillatory changes in intrinsic properties, including PIC amplitude and input conductance, consistent with over-correction rather than stable regulation [18]. In addition to intrinsic properties, inhibitory synapses onto motoneurons undergo significant remodeling in ALS, with changes in GABAergic and glycinergic receptor composition and synapse density reported at early disease stages [16], [19], [20], [21]. Whether these inhibitory changes are pathogenic drivers or compensatory responses to altered motoneuron activity remains unclear, as both reduced and enhanced inhibition could plausibly influence vulnerability [22], [23]. Systems-level analyses have proposed that ALS reflects instability of interacting control loops rather than failure of a single cellular process [24], [25].

Despite extensive research on changes to motoneuron excitability in ALS, few studies explicitly test whether ALS motoneurons respond differently than wild type motoneurons to the same chronic perturbation. In particular, genotype-specific differences in compensatory responses to sustained homeostatic challenges, such as chronic inhibition, remain largely unexplored at the level of intrinsic electrical properties. Diazepam is a benzodiazepine that exerts its effects by enhancing inhibitory neurotransmission mediated by GABA receptors. With chronic exposure, diazepam is known to induce compensatory weakening of inhibitory transmission and synaptic remodeling in multiple brain regions. Although motoneuron-specific data are limited, chronic benzodiazepine exposure provides a well-characterized inhibitory perturbation expected to engage homeostatic mechanisms rather than produce sustained suppression [26], [27]. In this study, we hypothesized that motoneurons in the mSOD1 mouse model of ALS exhibit excessive, high-gain homeostatic plasticity in response to chronic inhibitory perturbation compared to wild-type motoneurons. We predicted that such differences would manifest as genotype × treatment interaction effects, rather than as baseline genotype differences or uniform treatment effects. To test this hypothesis, we treated wild type and mSOD1 mice in the late pre-symptomatic phase with vehicle (sesame oil) or diazepam (15 mg/kg/day; for 10 days). On the last day of treatment, we made in vivo intracellular motoneuron recordings in anesthetized mice and analyzed motoneuron passive properties, active properties, and synaptic scaling of inhibitory inputs using mixed-effects statistical models. We found a significant genotype × treatment interaction effect for certain properties related to passive membrane integration and spike initiation: membrane time constant, peak input resistance, and recruitment current. This signified that mSOD1 motoneurons responded differently to the homeostatic challenge of chronic diazepam treatment. These findings provide evidence for high-gain homeostatic control of motoneuron firing output in mSOD1 motoneurons and reframes motoneuron excitability in ALS as a problem of altered feedback control and adaptive dynamics, rather than fixed hyper- or hypo-excitable states [18], [24].

## 2. Results

We postulate that in ALS, motoneurons have dysregulated homeostatic responses to aggregated proteins and other cellular stressors. We define excessive or high-gain homeostasis as a condition in which a sustained perturbation elicits disproportionately large compensatory changes in intrinsic properties in one genotype relative to another, despite comparable baseline properties and comparable exposure to the perturbation. This excessive homeostatic gain leads to homeostatic oscillations of electrical properties and ultimately motoneuron degeneration. In this study, we tested the hypothesis that sustained inhibition, a form of cellular stress, elicits over-correcting homeostatic plasticity in motoneurons of mSOD1 mice in the late pre-symptomatic stage.

### 2.1. Chronic treatment elevated diazepam and its active metabolites in the spinal cord

To induce a sustained inhibitory perturbation, we treated late pre-symptomatic mSOD1 mice and wild type littermates with diazepam (15 mg/kg/day) for 10 days. Diazepam and its active metabolites nordiazepam and oxazepam were elevated in lumbosacral spinal cord tissue of treated mice on the tenth day, detected using a validated liquid chromatography with tandem mass spectrometry (LC-MS/MS) method (Figure 1). Oxazepam was the predominant detected metabolite, followed by nordiazepam, while diazepam was present at the lowest levels. Temazepam was not detected (data not shown). For each detected compound, the Δ-Δ effect size between diazepam-treated wild type and diazepam-treated mSOD1 mice was negligible, indicating that the bioavailability of diazepam and its active metabolites was comparable for treated animals of both groups. Metabolite-to-parent ratios were consistent across treated animals (mean ≈2.9 and ≈8.2 for nordiazepam and oxazepam, respectively), indicating efficient metabolic conversion. In contrast, no analytes were detected above the lower limit of quantification in vehicle-treated groups, confirming method specificity and absence of background interference.

**Figure 1.**
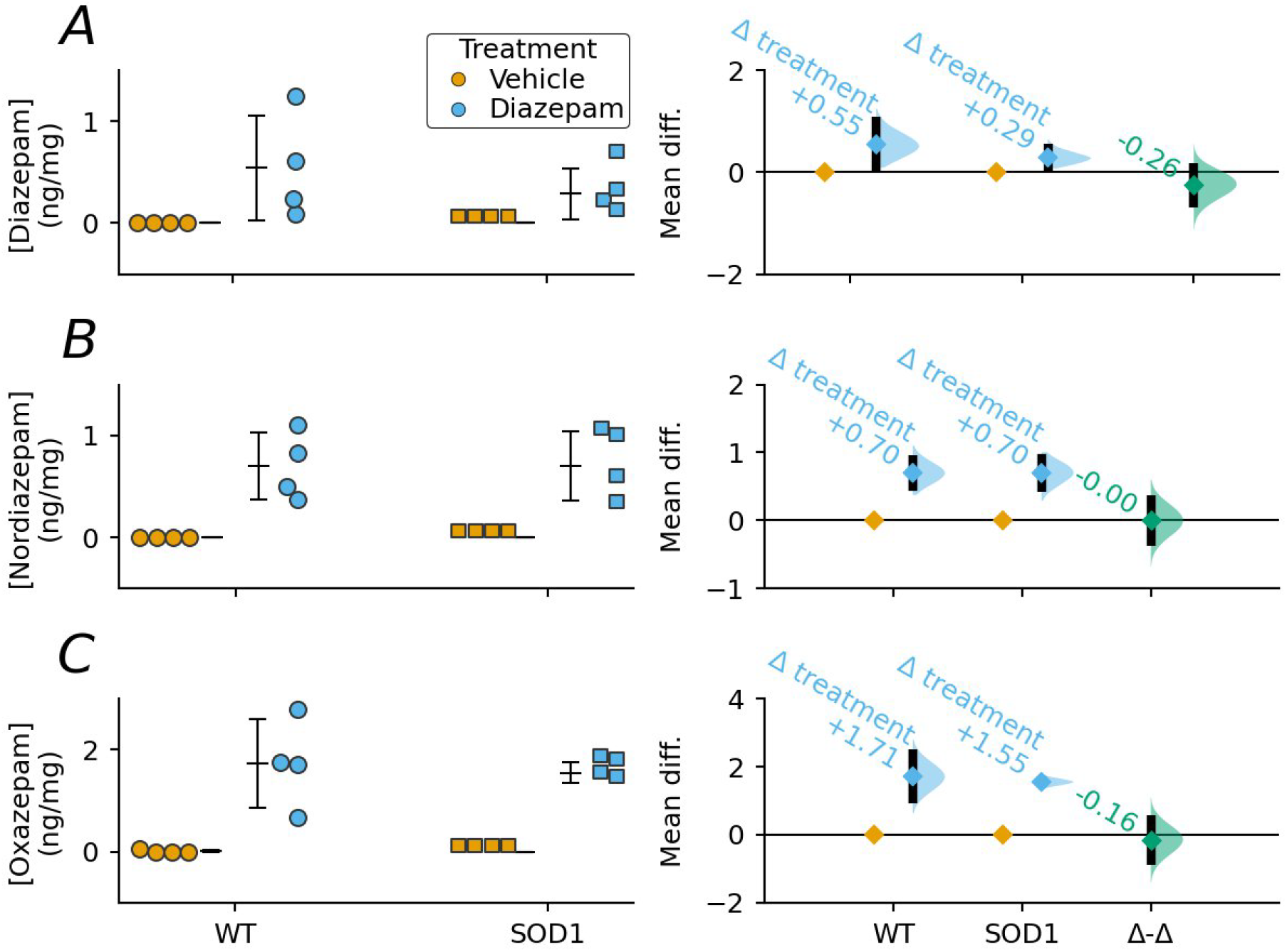
Cumming estimation plots of the level of diazepam and its active metabolites in the lumbosacral spinal cord of wild type (WT) and mSOD1 mice that were chronically treated with diazepam or vehicle (N=4 spinal cords per group). Scatterplots (left) and depictions of effect size (right) for (A) diazepam, (B) nordiazepam, and (C) oxazepam concentrations. In the scatterplots, each point represents a spinal cord and the lines next to the points show the mean ± standard deviation. In the depictions of effect size, the y-axis shows the bootstrapped distributions of the mean differences in concentration between diazepam-treated and vehicle-treated mice of the same genotype; on the far right, the y-axis shows the Δ-Δ effect size between diazepam-treated mSOD1 and diazepam-treated wild type mice; black lines represent 95% confidence intervals. Abbreviations: diff., difference.

### 2.2. Passive properties exhibited over-active homeostatic plasticity in mSOD1 motoneurons

On the tenth day of diazepam treatment, we performed in vivo intracellular motoneuron recordings in anesthetized wild type and mSOD1 mice and analyzed the effect of this inhibitory perturbation on motoneuron intrinsic properties (Figure 2). Overall, we performed intracellular recordings in 78 mice (40 WT and 38 mSOD1) and recorded 372 motoneurons. 96 motoneurons were recorded in WT mice injected with vehicle, 87 in WT mice injected with diazepam, 112 in mSOD1 mice injected with vehicle and 77 in mSOD1 mice injected with diazepam. Chronic diazepam treatment did not modulate resting membrane potential in wild type or mSOD1 motoneurons on the tenth day, as there were no significant genotype, treatment, or interaction effects detected using a linear mixed-effects model with Benjamini-Hochberg (BH) correction for multiple comparisons. On the other hand, statistical analysis of membrane time constant (τ_m_) revealed a significant main effect of genotype (FDR-corrected p-value q=0.0002) and a significant interaction between genotype and treatment (q=0.044). Group-level trends indicated that membrane time constant (τ_m_) increased preferentially in mSOD1 motoneurons under chronic diazepam, which is consistent with genotype-specific modulation of passive membrane integration. Sustained inhibition also induced a high-gain homeostatic response in peak input resistance. There was a significant main effect of genotype (q=0.048) and a significant genotype × treatment interaction effect (q=0.014). Like τ_m_, directional inspection of group means showed that peak input resistance increased selectively in SOD1 motoneurons following chronic diazepam treatment, with minimal changes in wild type motoneurons. The absence of change in wild type motoneurons on the tenth day of chronic treatment reflects that successful homeostasis was achieved in this group. Steady-state input resistance measured at the plateau showed no significant genotype, treatment, or interaction effects. Similarly, sustained inhibition did not induce high-gain homeostatic modulation of hyperpolarization-activated cation current (*I*_*h*_), as measured by the sag ratio [28]. Taken together, changes to τ_m_ and input resistance indicate a coordinated alteration of passive membrane properties specifically in mSOD1 motoneurons in response to chronic inhibition.

**Figure 2.**
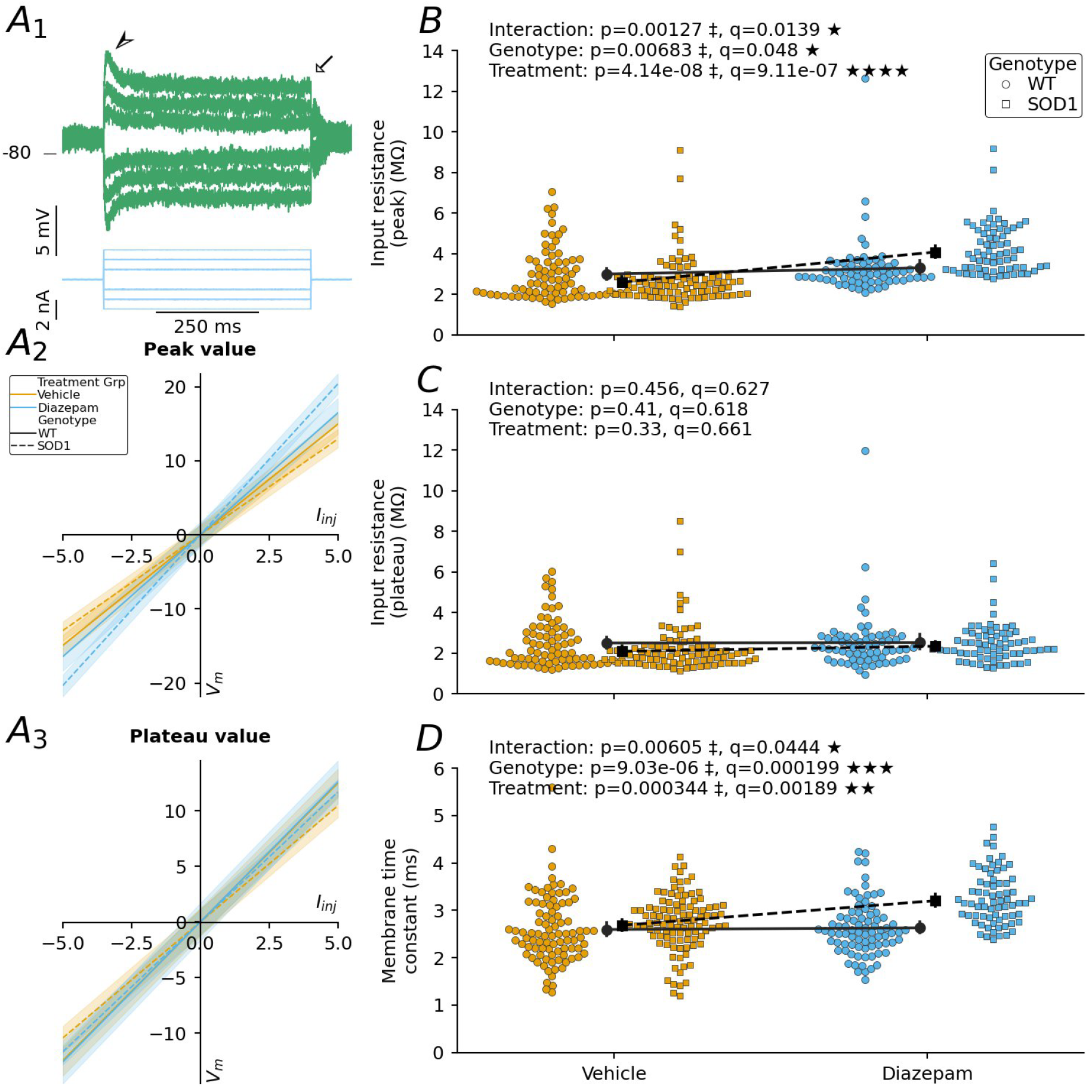
Modulation of motoneuron intrinsic electrical properties by chronic diazepam treatment. (**A**_**1**_) Input resistance was measured from peak (arrowhead) and plateau (arrow) voltage responses (top) to small-amplitude, 500-ms square current pulses (bottom). Average current-voltage relationships plotted from (**A**_**2**_) peak and (**A**_**3**_) plateau voltage responses for each of the four experimental groups: wild type (solid line) and mSOD1 (dashed line) motoneurons from vehicle-(orange) and diazepam-treated (blue) mice; shading surrounding each curve represents the 95% confidence interval. Scatterplots of motoneuron (**B**) peak input resistance, (**C**) plateau input resistance, and (**D**) membrane time constant (τ_m_). Wild type motoneurons are represented as circles and mSOD1 motoneurons are depicted as squares. Motoneurons from vehicle-treated mice are colored orange and those from diazepam-treated mice are colored blue. The difference in means between vehicle- and diazepam-treated mice of the same genotype is represented by a superimposed solid or dashed line for wild type motoneurons and mSOD1 motoneurons, respectively. *p* values correspond to the result of the mixed-effects statistical model before BH correction for multiple comparisons. *q* values correspond to the result after BH correction. *p*<0.05:^‡^; *q*<0.05: ★; *q*< 0.01: ★★; *q*<0.001: ★★★; *q*<0.0001: ★★★★.

### 2.3. Sustained inhibition caused high-gain homeostatic modulation of recruitment current in mSOD1 motoneurons

Because changes in input resistance and membrane time constant directly influence the current–voltage trajectory toward spike threshold, we next asked whether these passive adjustments translated into altered spike recruitment and firing behavior (Figure 3). We used triangular current ramps to characterize motoneuron firing behaviors. Diazepam treatment caused high-gain homeostatic modulation of recruitment current (Ion); Ion showed a significant main effect of genotype (q=0.0024) and a robust genotype × treatment interaction (q=0.002). Graphical inspection confirmed that chronic diazepam reduced Ion selectively in mSOD1 motoneurons, while wild type motoneurons showed little or no change in response to the chronic treatment. A related property, rheobase, which was measured using square current pulses of increasing amplitude, appeared similarly affected but most of these effects were no longer statistically significant after applying our standard BH correction for multiple comparisons. A significant main effect of genotype remained (q=0.048), signifying that rheobase was lower in pooled mSOD1 motoneurons; this outcome appears driven by a relatively smaller average rheobase in the diazepam-treated mSOD1 group. Overall, these results indicate a genotype-specific facilitation of spike initiation following sustained inhibitory perturbation. In contrast, firing gain and repetitive firing behaviors were not homeostatically modulated in wild type or mSOD1 motoneurons on the tenth day of chronic treatment. Specifically, current- and frequency-adaptation metrics (ΔI, the difference between derecruitment and recruitment currents; ΔF, the difference between offset and onset firing frequencies) and firing rate gain in the primary firing range did not have significant genotype, treatment, or interaction effects. These findings suggest that chronic diazepam treatment did not broadly alter steady-state firing gain or repetitive discharge properties in mSOD1 motoneurons, despite changes in recruitment and passive parameters unique to this group.

**Figure 3.**
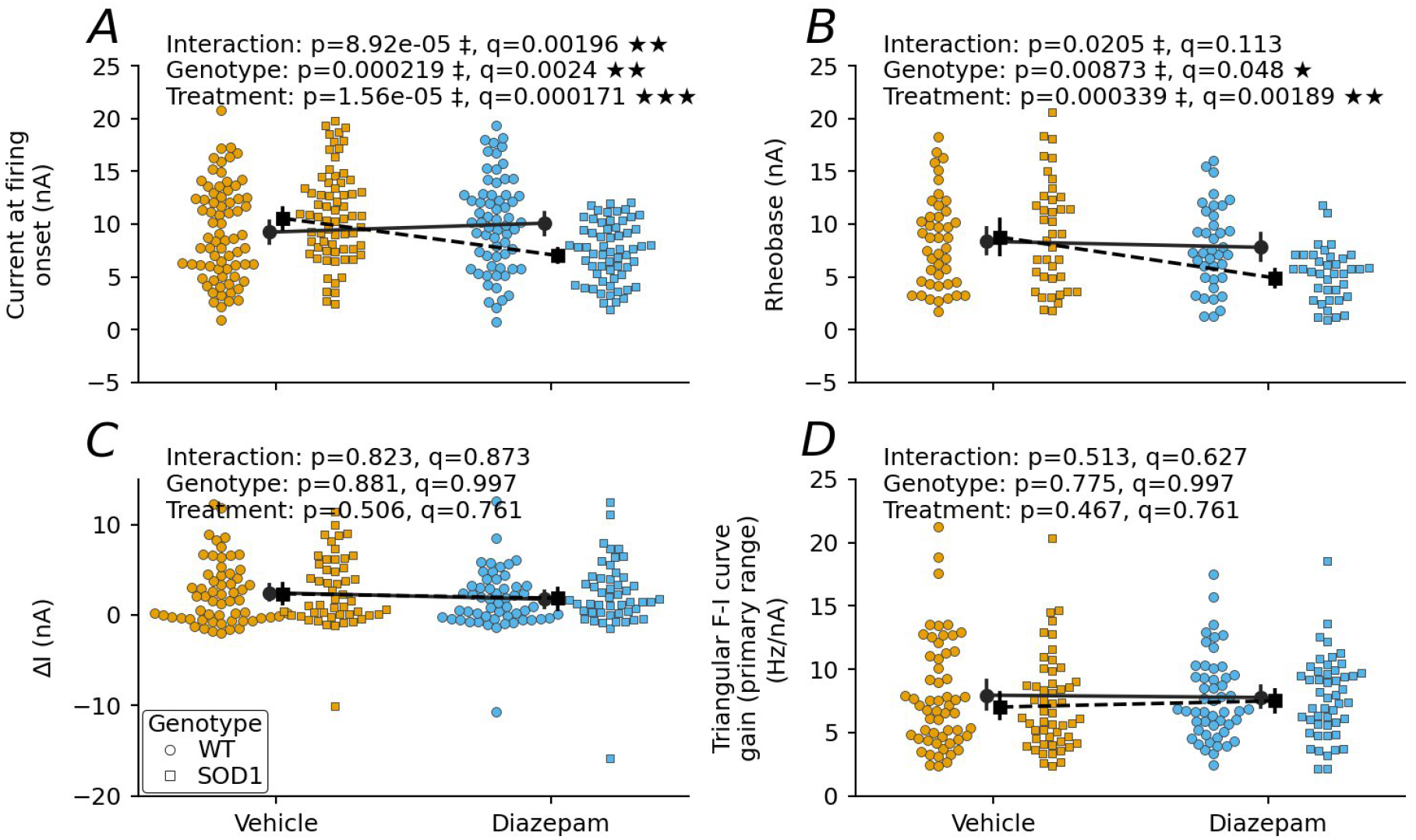
Effects of chronic diazepam treatment on motoneuron firing behavior. Scatterplots of **(A)** Ion (current at firing onset), **(B)** rheobase (current that 50%-of-the-time elicits an action potential), **(C)** ΔI (the difference between derecruitment and recruitment currents), and **(D)** primary firing range gain. Wild type motoneurons are represented as circles and mSOD1 motoneurons are depicted as squares. Motoneurons from vehicle-treated mice are colored orange and those from diazepam-treated mice are colored blue. The difference in means between vehicle- and diazepam-treated mice of the same genotype is represented by a superimposed solid or dashed line for wild type motoneurons and mSOD1 motoneurons, respectively. *p* values correspond to the result of the mixed-effects statistical model before BH correction for multiple comparisons. *q* values correspond to the result after BH correction. *p*<0.05:^‡^ ; *q*<0.05: ★; *q*<0.01: ★★; *q*<0.001: ★★★.

To test whether the exaggerated homeostatic response to chronic diazepam treatment included changes to spike properties, we analyzed action potential waveforms from averaged single spikes elicited by brief current pulses. There were no significant genotype, treatment, or interaction effects for the properties of spike height, spike overshoot, and spike width. Afterhyperpolarization (AHP) properties, including amplitude, time-to-peak, and half-relaxation time, also showed no significant genotype, treatment, or interaction effects. Thus, spike shape and AHP dynamics remained stable across genotypes and treatments. A complete list of electrophysiological properties that we evaluated in this study can be found in Table 1.

**Table 1.**
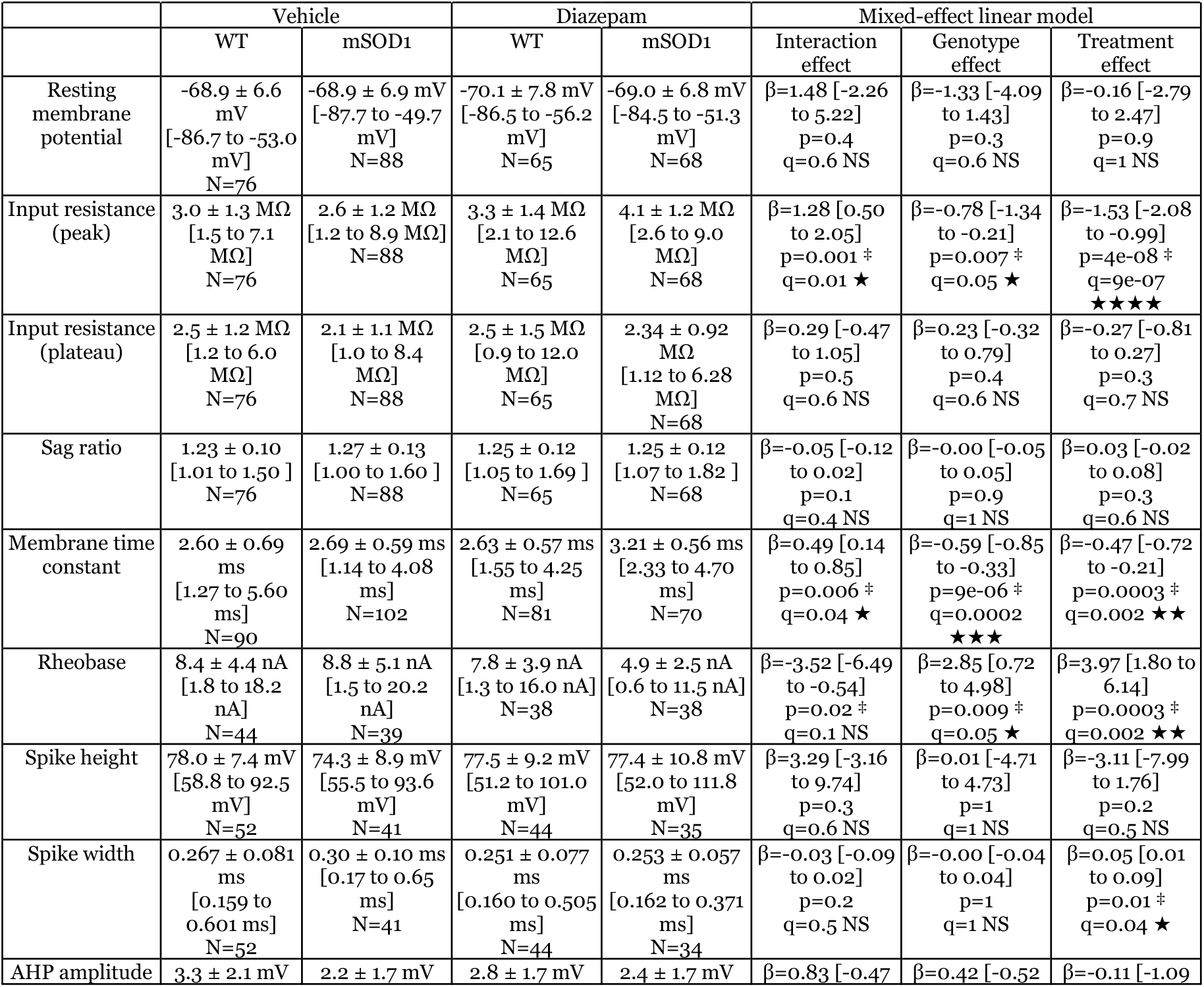
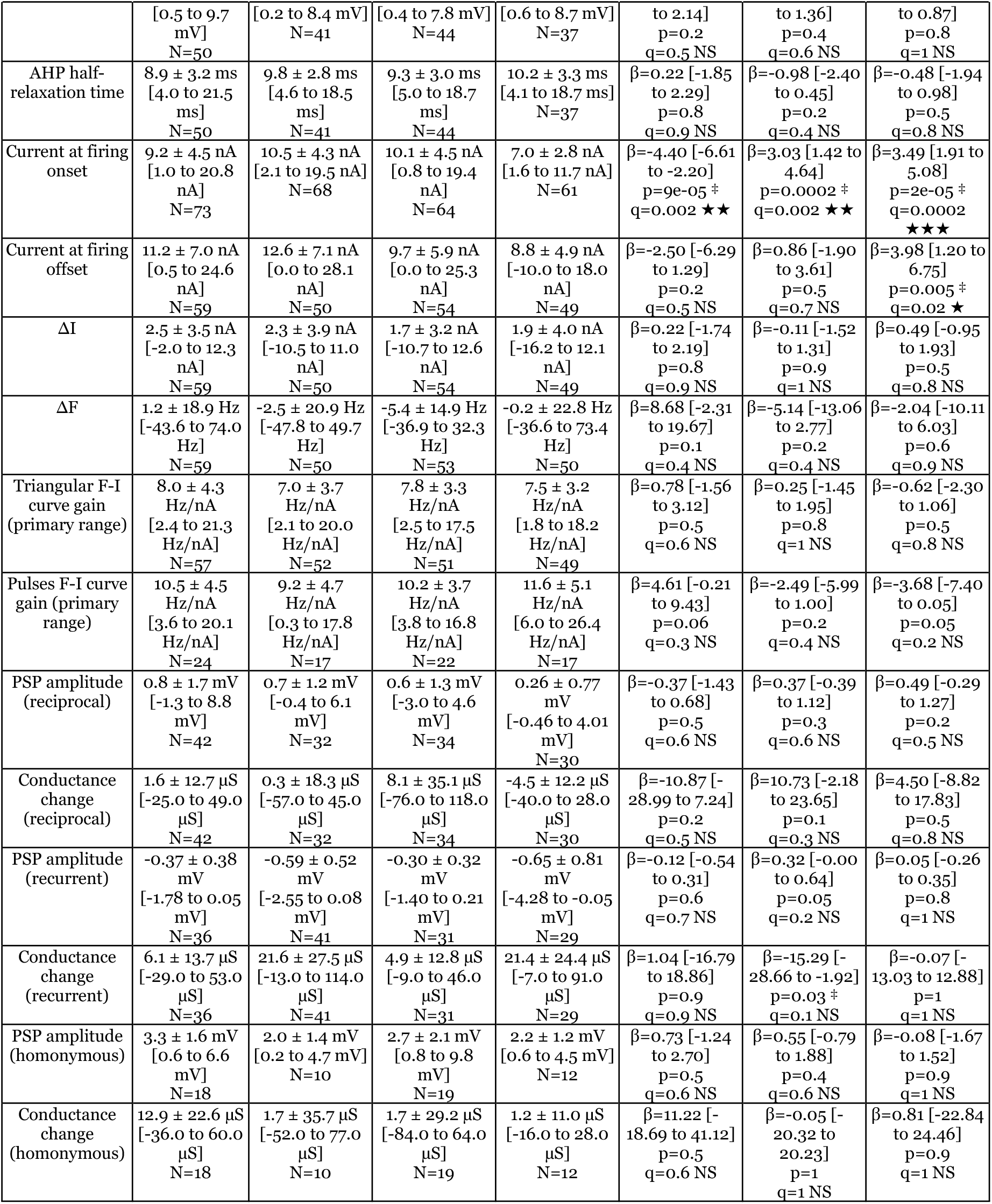
Summary of electrophysiological properties measured in this study. For each property, the mean ± standard deviation is reported for each group (left columns); range is provided in brackets; number of cells (N) is listed beneath the range. Results from mixed-effects linear model statistical testing are presented as follows (right columns): value of the factor β for each effect, with its 95% confidence interval; *p* values correspond to the result of the mixed-effects statistical model before BH correction for multiple comparisons. *q* values correspond to the result after BH correction. *p*< 0.05:^‡^ ; *q*<0.05: ★; *q*<0.01: ★★; *q*<0.001: ★★★. Abbreviations: AHP, after-hyperpolarization; ΔF, difference between offset and onset firing frequencies; F-I, frequency-current; ΔI, difference between derecruitment and recruitment currents; PSP, post-synaptic potential; WT, wild type.

### 2.4. Diazepam treatment did not elicit homeostatic synaptic scaling of synaptic inputs

We next asked whether exaggerated homeostatic responses in mSOD1 motoneurons extended beyond intrinsic properties to synaptic inputs, which would indicate a more global homeostatic dysregulation. To maintain normal firing rates despite chronically reduced or elevated network activity, neurons globally scale up or down synaptic strength through a post-synaptic mechanism called synaptic scaling [29], [30], [31]. We hypothesized that sustained inhibition in the spinal cord would induce synaptic scaling of two major synaptic pathways onto motoneurons, reciprocal and recurrent inhibition [32].

#### 2.4.a. Reciprocal connections

Experiments performed in cats have shown that stimulation of Ia afferent fibers from one muscle inhibits the antagonist motor pool through inhibitory interneurons called the Ia inhibitory interneurons [32], [33], [34]. This pathway, called reciprocal inhibition, is thought to facilitate alternation between flexor and extensor muscles that act at the same joint during locomotion, such as the tibialis anterior and extensor digitorum longus (TA and EDL, respectively; ankle flexors; innervated by the deep peroneal nerve) and triceps surae (TS; ankle extensor; innervated by the TS nerve). In our preparation, we investigated this pathway by recording intracellularly from TS motoneurons while electrically stimulating the deep peroneal (DP) nerve (or vice-versa) at twice the threshold intensity of the group I fibers (Figure 4A_1_). Because the inhibitory post-synaptic potential (PSP) can be very small when the membrane potential is close to the chloride reversal potential, we systematically depolarized the membrane potential using a bias current to uncover any IPSP that could be too small to observe at rest. While in some motoneurons we did observe that the stimulation of the DP nerve elicited purely inhibitory PSPs on TS motoneurons, we were surprised to observe that other motoneurons received purely excitatory PSPs, mixed responses, or no responses at all (Figure 4C). Our observation of reciprocal excitation appears to highlight an interesting species-specific difference in reciprocal synaptic connections between motor pools in the mouse. We tested whether the proportion of motoneurons exhibiting these different types of responses was different between our experimental groups. A multinomial mixed-effects model revealed a genotype × treatment interaction. This interaction was most strongly felt in the prevalence of EPSPs (posterior estimate = −10.27, 95% CI −21.28 to −1.38), indicating that treatment effects on EPSP probability differ between genotypes. Evidence for interaction effects on other types of responses was inconclusive. Overall, mSOD1 motoneurons had a smaller proportion of EPSPs after diazepam treatment than wild type motoneurons.

**Figure 4.**
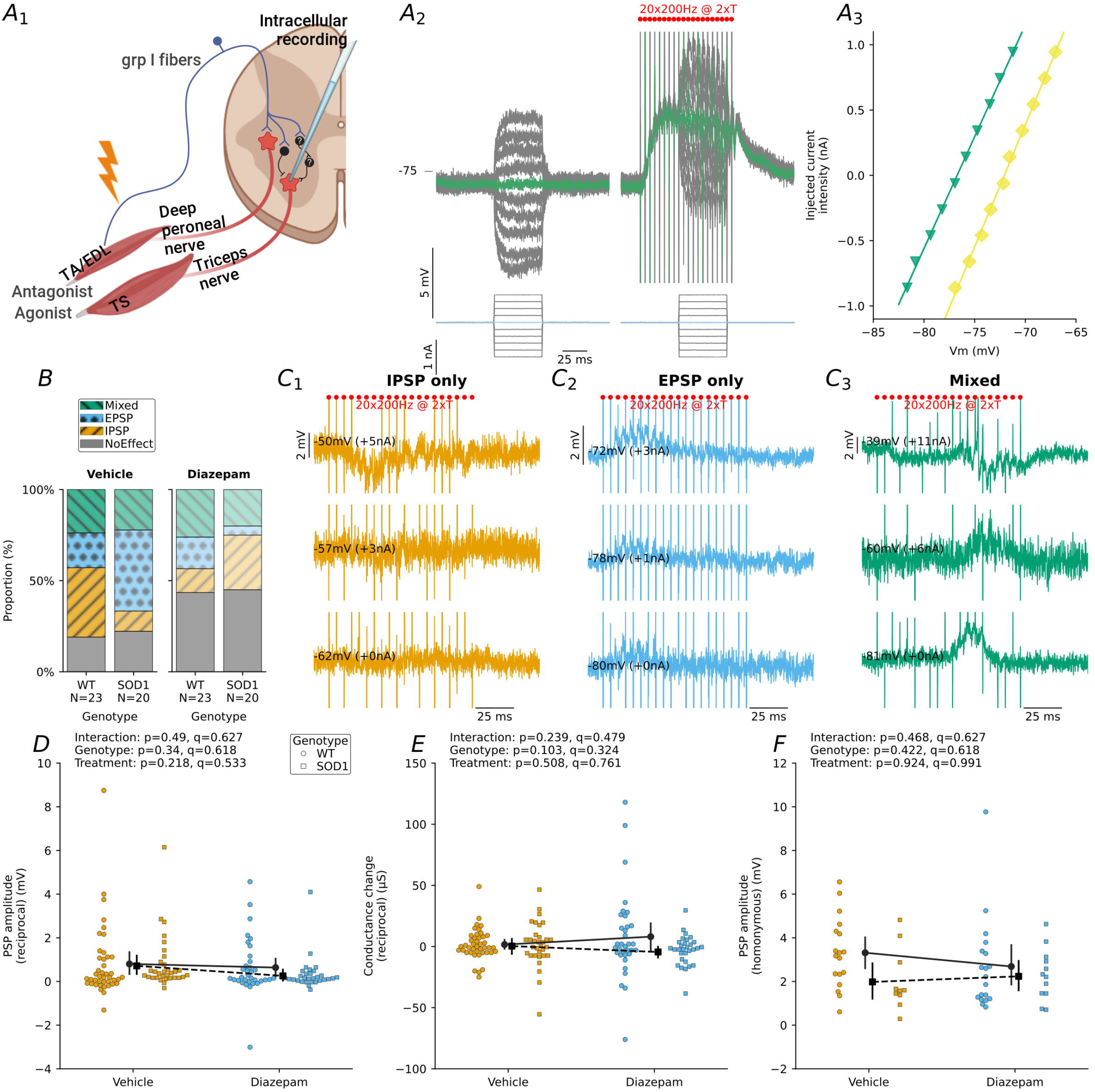
Effects of chronic inhibition by diazepam on the strength of reciprocal synaptic inputs onto motoneurons. **(A_1_)** Schematic of the preparation for *in vivo* recording of reciprocal connections in motoneurons. **(A_2_)** Examples of voltage responses to 50-ms current pulses applied 300-ms prior to antagonist nerve stimulation (pre-stim) and a second series of pulses 40-ms after the first shock in antagonist nerve stimulation (a series of 20 shocks at 200 Hz; mid-stim). For the pre-stim and mid-stim series, the steps in which no current is passed is colored green. **(A_3_)** I-V relationships between current pulse amplitudes and membrane potentials recorded pre- and mid-antagonist-nerve stimulation. **(B)** Stacked bar charts depicting the relative frequencies that **(C_1_)** IPSPs, **(C_2_)** EPSPs, **(C_3_)** mixed, or no responses were observed in motoneurons in response to antagonist nerve stimulation (20 x 200 Hz at 2 x T). These recordings used a bias current to depolarize the membrane potential above the chloride reversal potential; this controlled direction (inward) and augmented amplitude of IPSPs when present. **(D)** Scatterplots of peak PSP amplitude measured mid-antagonist-nerve stimulation in the absence of injected current (as in green trace on the right in A_2_). **(E)** Synaptic conductance changes during steady-state reciprocal inhibition, estimated from the difference in slopes of the I-V relationships depicted in A_3_. **(F)** Scatterplots of peak PSP amplitude evoked mid-homonymous-nerve stimulation in the absence of injected current. Wild type motoneurons are represented as circles and mSOD1 motoneurons are depicted as squares. Motoneurons from vehicle-treated mice are colored orange and those from diazepam-treated mice are colored blue. The difference in means between vehicle- and diazepam-treated mice of the same genotype is represented by a superimposed solid or dashed line for wild type motoneurons and mSOD1 motoneurons, respectively. *p* values correspond to the result of the mixed-effects statistical model before BH correction for multiple comparisons. *q* values correspond to the result after BH correction. Abbreviations: EDL, extensor digitorum longus; EPSP, excitatory post-synaptic potential; grp, group; IPSP, inhibitory post-synaptic potential; PSP, post-synaptic potential; T, threshold; TA, tibialis anterior; TS, triceps surae; WT, wild type.

**Figure 5.**
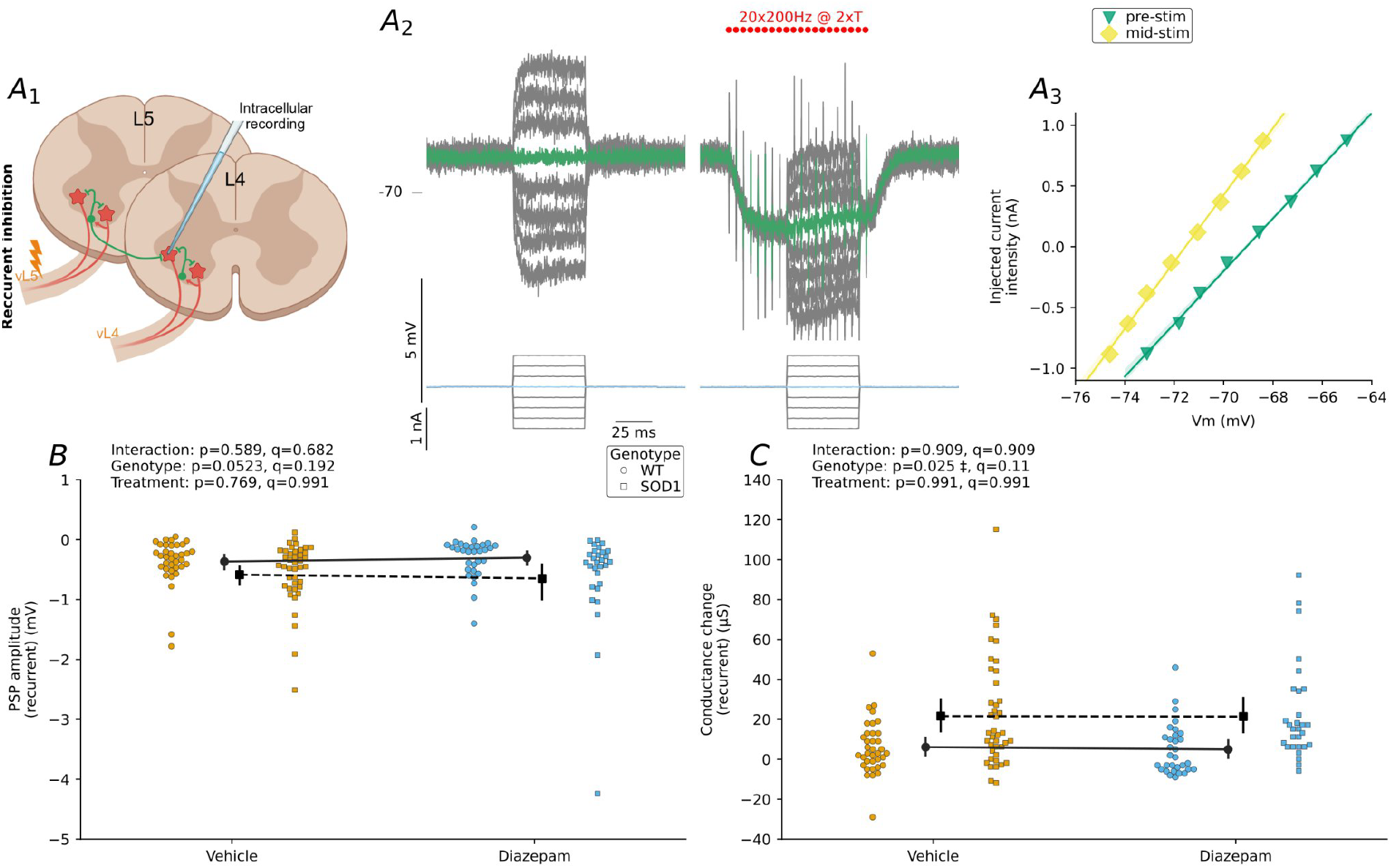
Effects of chronic inhibition by diazepam on the strength of recurrent inhibition of motoneurons. (A_1_) Schematic of the preparation for *in vivo* recording of recurrent inhibition in L4 motoneurons. (A_2_) Examples of voltage responses to 50-ms current pulses applied 300-ms prior to adjacent ventral root stimulation (pre-stim) and a second series of pulses 40-ms after the first shock in adjacent ventral root stimulation (a series of 20 shocks at 200 Hz; mid-stim). (A_3_) I-V relationships analogous to those described in A_3_. **(B)** Scatterplots of peak PSP amplitude evoked during the recurrent inhibition protocol (measured on the response where no current was injected, highlighted in green in A_2_). **(C)** Scatterplots of change in synaptic conductance during steady-state recurrent inhibition estimated from the difference in slopes of the I-V curves in A_3_. Wild type motoneurons are represented as circles and mSOD1 motoneurons are depicted as squares. Motoneurons from vehicle-treated mice are colored orange and those from diazepam-treated mice are colored blue. The difference in means between vehicle- and diazepam-treated mice of the same genotype is represented by a superimposed solid or dashed line for wild type motoneurons and mSOD1 motoneurons, respectively. *p* values correspond to the result of the mixed-effects statistical model before BH correction for multiple comparisons. *q* values correspond to the result after BH correction. Abbreviations: PSP, post-synaptic potential; T, threshold; WT, wild type.

We then measured peak PSP amplitude at RMP during antagonist nerve stimulation, but noted that the magnitude and direction were highly variable (Figure 4D), dependent on how close the RMP was to the chloride reversal potential. To avoid this complication, we instead measured the change in conductance associated with synapse activation as a proxy for reciprocal synaptic strength. We measured synaptic conductance during steady-state reciprocal inhibition (or homonymous excitation) using a current-voltage (I-V) curve generated from voltage responses evoked by a series of square current pulses before and during nerve stimulation (Figure 4A and E). Measures of reciprocal PSPs and conductance did not show significant genotype, treatment, or genotype × treatment interaction when analyzed using a linear mixed-effects model with BH correction for multiple comparisons. This result indicates that reciprocal connections input strength was not detectably modified by chronic diazepam in a genotype-specific manner.

We also elicited homonymous excitation by electrically stimulating the TS nerve (or vice versa) while using a negative bias current to block the antidromic spike. Similarly, we found that measures of homonymous excitatory PSPs and synaptic conductance did not exhibit significant main or interaction effects. These results demonstrate that excitatory synaptic drive onto motoneurons was not detectably altered by chronic diazepam in either genotype.

#### 2.4.b. Recurrent inhibition

Since the previous circuit proved to not be exclusively inhibitory, we then focused on the recurrent inhibition circuit. Recurrent inhibition is a negative feedback loop in which motoneuron axon collaterals activate Renshaw cells, which inhibit homonymous motoneurons and/or heteronymous motoneurons innervating a synergistic muscle [32]. We tested whether chronic diazepam treatment induced homeostatic synaptic scaling of recurrent inhibition. These experiments were performed in a different set of animals, in which the ventral roots of two adjacent spinal segments were cut distally and mounted on bipolar stimulation electrodes. We evoked recurrent inhibition by electrical stimulation of the ventral root adjacent to that which the recorded motoneuron belonged to [22], [35]. We did not observe homeostatic synaptic scaling of recurrent inhibition, as there were no significant main treatment or genotype × treatment interaction effects for recurrent inhibitory PSP or conductance properties (Figure 4).

Overall, we did not detect homeostatic modulation of reciprocal or recurrent synaptic connections in response to chronic diazepam treatment. The absence of synaptic scaling suggests that excessive homeostatic gain in mSOD1 motoneurons is not uniform and is selectively affecting intrinsic electrophysiological properties.

## 3. Discussion

In this study, we used in vivo electrophysiology to test whether motoneurons from pre-symptomatic mSOD1 mice modulate excitability differently than wild type motoneurons in response to the same chronic inhibitory perturbation. The sustained inhibitory perturbation that we implemented in this study was chronic treatment with diazepam, a benzodiazepine that enhances inhibitory neurotransmission mediated by GABA receptors.

We performed sharp electrode intracellular motoneuron recordings in anesthetized mice to determine if chronic inhibition by diazepam modulated motoneuron intrinsic properties, firing behaviors, and synaptic inputs differently in mSOD1 mice compared to their wild type littermates. This outcome would be signified by a statistically significant genotype × treatment interaction effect when using mixed-effects statistical models and BH correction for multiple hypothesis tests. Implementing this conservative method, significant interaction effects were confined to a small subset of intrinsic properties. Specifically, we found that RinPeak, τ_m_, and recruitment current (Ion) exhibited genotype-specific responses to chronic diazepam, all related to passive membrane integration and spike initiation. In contrast, firing gain, spike waveform properties, and synaptic inputs remained largely unaffected by chronic diazepam treatment. This pattern indicates that chronic inhibitory perturbation selectively triggers overactive intrinsic compensatory mechanisms in mSOD1 motoneurons rather than producing widespread changes in firing or synaptic transmission. The selectivity of the effects supports a selective engagement of intrinsic compensatory mechanisms, rather than a generalized destabilisation of motoneuron electrophysiological properties.

We measured synaptic scaling of reciprocal inputs onto motoneurons after sustained inhibition by diazepam, but found evidence that these reciprocal Ia synaptic connections are not purely inhibitory in the mouse. This starkly contrasts what is seen in the cat [36], [37], but is consistent with prior observations made by Horstman et al. (2019), who found that conditioning stretches of the TA sometimes facilitated the stretch reflex in the rat gastrocnemius [38]. Future studies are needed to define the differences between cats and rodents in reciprocal connections between agonist and antagonist motor pools.

We selected our dosing strategy because former studies have demonstrated that it avoids daily withdrawal and results in chronic receptor occupancy in the mouse forebrain (still 34% occupancy at 24-h post-injection; [39], [40]). Our treatment method (diazepam in sesame oil; 15 mg/kg/day; SID SQ for 10 days) led to the detection of diazepam, nordiazepam, and oxazepam in the lumbosacral spinal cord of treated wild type and mSOD1 mice. The concentrations of each detected compound were comparable between diazepam-treated wild type and diazepam-treated mSOD1 spinal cords. This demonstrates that the larger homeostatic response of motoneurons from diazepam-treated mSOD1 mice to chronic diazepam treatment was not attributable to greater bioavailability of diazepam and/or its active metabolites. Instead, the greater modulation of recruitment and passive properties in mSOD1 motoneurons likely reflects dysregulated homeostatic plasticity.

Our finding of high-gain homeostatic plasticity in mSOD1 motoneurons is consistent with the few other studies that have directly tested whether homeostatic responses of wild type and mSOD1 motoneurons are different or not. Kuo et al. (2020)[17] perturbed anabolism in neonatal wild type and mSOD1 mice by chronic (7-days) treatment with PF-47, an inhibitor of the target of rapamycin (mTOR) signaling pathway. Like us, they found that by the day of electrophysiological recording (using whole-cell patch clamp), drug-treated wild type motoneurons exhibited normal electrophysiological properties (reflecting successful homeostasis) while drug-treated mSOD1 motoneurons showed decreased soma size, conductance and PICs (reflecting excessive homeostasis to the same perturbation). Similarly, Schuster et al. (2012)[41] perturbed excitability in embryonic wild type and mSOD1 mouse primary spinal cord cultures by chronic (4-9 days) treatment with riluzole. They observed a similar result by whole-cell patch clamp, that riluzole decreased PIC amplitude in riluzole-treated mSOD1 but not wild type motoneurons. Recent work by Mahrous et al. (2026) tested whether chronic treatment (10-days) of pre-symptomatic mSOD1 mice with riluzole was ineffective to normalize motoneuron excitability because of over-correcting homeostatic responses in motoneurons [42]. Consistent with our results, chronic treatment with riluzole altered PICs and firing behavior in mSOD1 but not wild type motoneurons recorded in an ex vivo sacral cord preparation. Here, we provide the first in vivo evidence for hypervigilant homeostasis in motoneurons of adult mSOD1 mice during the pre-symptomatic period. Antonucci et al. (2026)[43] treated pre-symptomatic wild type and mSOD1 mice with β-adrenergic receptor (Adrb2/Adrb3) agonists. Acute treatment increased excitability of both wild type and mSOD1 mice, but after chronic treatment (10-days), no electrophysiological properties showed significant genotype × treatment interaction effects - this suggests that both wild type and mSOD1 motoneurons adapted successfully to this particular sustained perturbation. Thus, it appears that not all homeostatic feedback loops are equally dysregulated in ALS. While cAMP/PKA-mediated neuromodulatory loops appear capable of successful long-term adaptation, inhibitory perturbations engaging GABAergic signaling preferentially experience excessive intrinsic compensation.

In conclusion, this study provides evidence for over-correcting homeostatic control of motoneuron firing output in mSOD1 mice during the late pre-symptomatic stage. While wild type motoneurons executed successful homeostasis of electrical properties by ten days of chronic inhibition, recruitment and passive parameters of mSOD1 motoneurons were abnormal, signifying high-gain homeostatic plasticity. Our work recontextualizes altered motoneuron excitability in ALS as a problem of dysregulated feedback control and homeostatic dynamics, rather than fixed pathological states of excitability.

## 4. Materials and Methods

### 4.1. Ethical statement

All experimental procedures conducted with animals were approved by the University of Rhode Island’s Institutional Animal Care and Use Committee and performed in accordance with the National Institutes of Health Guide for the Care and Use of Laboratory Animals.

### 4.2. Animal housing and husbandry

B6SJL-Tg(SOD1*G93A)1Gur/J mice were obtained from The Jackson Laboratory (JAX○ stock no. 002726) and maintained in our colony by breeding with non-transgenic B6SJLF1/J mice (JAX○ stock no. 100012). Mice were housed in a controlled vivarium with a 12:12 h light-dark photoperiod, were monitored for health, and had access to food and water ad libitum. mSOD1 and non-transgenic littermates (wild type) were used for in vivo electrophysiology during the late pre-symptomatic stage, aged between 45 and 55 days old (wild type: 49.6 ± 2.0 days old, N=40; mSOD1: 49.4 ± 1.4 days old, N=38).

### 4.3. Chronic diazepam treatment

To induce a sustained inhibitory perturbation in motoneurons, we treated wild type and mSOD1 mice in the pre-symptomatic phase with vehicle (sesame oil; Sigma, cat. # S3547) or diazepam (15 mg/kg/day; Sigma, cat. # D0899; suspended by sonication in sesame oil) delivered subcutaneously once daily for 10 days, with the electrophysiology experiment performed on the 10th day of injection. We used a 31G needle to inject the intended volume (+20%) under the skin between the shoulders, which minimized backflow and compensated for the estimated leakage.

### 4.4. Surgical procedure

The surgical procedure has been previously described [28], [44], [45]. Briefly, pre-medications atropine (0.25 mg/kg; Covetrus, cat. # 074760) and dexamethasone (2 mg/kg; Covetrus, cat. # 070789) were administered subcutaneously to prevent salivation and edema during the surgery. Ten minutes later, anesthesia was induced by intraperitoneal injection of a cocktail containing fentanyl (0.025 mg/kg; Covetrus, cat.# 071169), midazolam (7.5 mg/kg; Covetrus, cat.# 072622) and dexmedetomidine (0.5 mg/kg; Covetrus, cat.# 034362). Core temperature was measured using a rectal thermometer and kept between 36.5 °C and 37.5 °C using an infrared heating lamp and electric heating pad; heart rate was measured by electrocardiogram and an adequate surgical plane was confirmed by a stable heart rate of 400-500 beats per minute and absence of response to toe pinch. A tracheostomy was performed and the mouse was placed under artificial respiration with pure oxygen (SAR-1000 ventilator; CWE, Ardmore, PA). The ventilator settings were monitored throughout the procedure and adjusted to maintain end-tidal PCO2 at ∼4% (Micro-Capstar; CWE); an adequate surgical plane was confirmed by stable end-tidal PCO2. Catheters were inserted into the left and right external jugular veins; one administered maintenance doses of anesthesia (1.25 µg/kg fentanyl; 0.375 mg/kg midazolam; 0.025 mg/kg dexmedetomidine; delivered as needed) and the other was used for continuous perfusion with HydraVol IV○ (100 µL/h; Covetrus, cat.# 086328). Two pairs of horizontal bars (Cunningham Spinal Adaptor;

Stoelting, Dublin, Ireland) were applied underneath the T13 and L2 transverse processes to immobilize the vertebral column; a suspended clamp on a sacral spinous process immobilized the sacrum. L3-S1 spinal segments were exposed by dorsal laminectomy. A custom-made plastic ring was secured around the exposed spinal cord and silicon sealant (Body Double Fast Set, Smooth-On Inc., Macungie, PA) was applied to create a recording chamber. The chamber was filled with mineral oil to prevent the spinal cord from drying out.

The surgeries were performed to enable recordings of recurrent or reciprocal inhibition in motoneurons. For experiments that included testing of recurrent inhibition, adjacent dorsal roots (L3-L4 or L4-L5) were transected proximal to the spinal cord to eliminate sensory-driven synaptic input, and the corresponding ventral roots were cut distally and mounted on bipolar stimulation electrodes. For experiments that included testing of reciprocal inhibition, the hamstring, sural, and all branches of the tibial nerve were cut except the TS nerve, which was mounted on a monopolar stimulation electrode. All branches of the common peroneal nerve were resected except the DP nerve, which was transected distally and mounted on a bipolar stimulation electrode. Muscle and nerve were covered in a mixture of petroleum jelly and mineral oil to prevent desiccation. To improve the stability of the intracellular recordings, respiratory movements were blocked by bolus intravenous administration of the paralytic pancuronium bromide (0.20 mg/kg; Sigma, cat. # P1918).

### 4.5. Electrophysiological recordings

Intracellular recordings were made using glass microelectrodes (tip diameter: 1.0–1.5 µm; resistance: 20-35 MΩ) filled with 2 M potassium acetate (Sigma, cat.# P1190). Intracellular recordings were obtained with an Axoclamp 2B amplifier (Molecular Devices, San Jose, CA, USA) and Spike2 software (CED, Cambridge, UK). All recordings were obtained using discontinuous current clamp (8 kHz; [46]) and sampled at 30 kHz.

For experiments that tested homeostatic synaptic scaling of recurrent inhibition in motoneurons belonging to L3-L5 spinal segments, each impaled motoneuron was classified by the spinal segment to which it belonged, which was determined by ventral root stimulation and the presence of an antidromic action potential. For the reciprocal inhibition experiments, each impaled motoneuron was classified by the motor pool to which it belonged, which was determined by TS or DP nerve stimulation and the presence of an antidromic action potential.

In both types of experiments, motoneuron input resistance was measured from peak and plateau voltage responses evoked by a series of small-amplitude 500-ms square current pulses [28]. Membrane time constant was measured using the ‘peeling’ method on the relaxation of the membrane potential after 1-ms hyperpolarizing current pulses [47]. Triangular current ramps (0.5-2 nA/s) were used to evaluate motoneuron firing properties. Voltage threshold for action potential firing was measured for the first spike evoked on the ramp, and was defined as the point where the first derivative of the voltage reached 10 mV/ms [48]. Recruitment current (Ion) was defined as the current intensity at voltage threshold; derecruitment current (Ioff) was defined as the current intensity at the time of the last spike on the descending ramp; the difference between derecruitment and recruitment currents (ΔI) was calculated. The frequencies of the first inter-spike interval on the ascending ramp and the last inter-spike interval on the descending ramp were used to calculate the difference between offset and onset firing frequencies (ΔF). Instantaneous firing frequency was plotted against injected current intensity at the time of the spike to characterize the frequency-current (F-I) relationship. To estimate gain, the slope of the F-I curve in the linear primary range, when present, was measured by linear regression. Primary firing range gain was also measured from F-I curves generated by injection of 500-ms depolarizing square current pulses. Spike properties (e.g., spike height; spike width at half-amplitude; AHP amplitude, measured as the potential difference between the trough of the medium AHP and resting membrane potential; AHP half-relaxation time, measured as the duration of time it takes for the medium AHP to relax to half its peak amplitude) were calculated from an average of ∼5-15 spikes elicited by current pulses of 1-ms duration repeated at a frequency of 3.3 Hz; a bias current was used to bring the resting membrane potential to -60 mV. Rheobase, defined as the current intensity required to elicit an action potential 50% of the time, was measured in response to 20-ms current pulses of increasing amplitude.

We measured recurrent inhibition as described previously [22]. Briefly, the ventral root adjacent to that which the impaled motoneuron belonged was stimulated at an intensity that elicited supramaximal recurrent inhibition. The presence of inhibition was visually confirmed by simultaneous injection of depolarizing bias current, which increased the driving force for chloride influx and therefore the discernibility of the inhibitory post-synaptic potentials. To measure inhibitory post-synaptic potential amplitude and estimate synaptic conductance during steady-state recurrent inhibition, we recorded a series of small amplitude 50-ms current pulses applied 300-ms prior to adjacent ventral root stimulation (control series) and a second series of pulses 40-ms after the first shock in the ventral root stimulation (a series of 20 shocks at 200 Hz; test series). The membrane potential in response to each current pulse was averaged from ∼5-10 sweeps. The change in synaptic conductance during steady state recurrent inhibition was measured from the difference in slopes of I-V curves generated by plotting current pulse amplitude versus the change in membrane potential for the control and test series. The peak PSP amplitude was measured using the step in the test series in which no current was injected.

To elicit reciprocal inhibition, the nerve to the antagonist muscle (TS or DP nerve) was stimulated at approximately twice the threshold intensity of the group I afferent volley, which was detected using a silver ball electrode positioned on the dorsal surface of the spinal cord. Reciprocal inhibition was visually distinguished from reciprocal excitation by simultaneous injection of depolarizing bias current which increased the driving force for chloride influx and therefore the discrimination of the inhibitory PSPs from excitatory PSPs. PSP amplitude and estimated synaptic conductance during steady state reciprocal inhibition was measured as described above.

At the end of the experiment, the anesthetized mouse was euthanized with an overdose of pentobarbital. The lumbosacral spinal cord was dissected from the vertebral column, rinsed in phosphate buffered saline, blotted on sterile gauze, and frozen at -80 °C until mass spectrometry.

### 4.6. Liquid chromatography with tandem mass spectrometry (LC-MS/MS)

Spinal cord tissue samples were retrieved from frozen storage (−80 °C) and accurately weighed using a precision balance. Each sample was homogenized using a handheld homogenizer in ice-cold 1% (v/v) formic acid to produce a 6-fold (w/v) homogenate. For protein precipitation, 5 µL of homogenate was mixed with 95 µL of acetonitrile. The mixture was vortexed vigorously for 30 seconds and centrifuged at 10,000 × g for 15 minutes at 4 °C. The supernatant was diluted with 200 µL of ultrapure water, filtered through a 0.2 µm syringe filter, and the resulting clarified filtrate was injected into the LC-MS/MS system for quantitative analysis.

Analyte separation was performed on an Agilent 1260 Infinity II LC system equipped with a Waters Acquity BEH C18 column (50 × 2.1 mm, 1.7 µm; Waters). The column temperature was maintained at 45 °C, and analytes were eluted at a flow rate of 0.5 mL/min using a binary gradient of 0.1% formic acid in water (solvent A) and 0.1% formic acid in acetonitrile (solvent B). The gradient program was as follows: 60% A/40% B at 0.2 min, 50% A/50% B at 1.0 min, 0% A/100% B from 2.1 to 4.4 min, and re-equilibration to 90% A/10% B from 4.5 to 5.5 min.

Detection was carried out using an Agilent 6470 triple quadrupole mass spectrometer operating in positive ion mode with Agilent Jet Stream electrospray ionization (AJS-ESI). Source parameters were optimized as follows: capillary voltage, 2500 V; nozzle voltage, 500 V; drying gas temperature, 325 °C; drying gas flow, 13 L/min; nebulizer pressure, 40 psi; sheath gas temperature, 375 °C; and sheath gas flow, 11 L/min. Analytes were monitored using dynamic multiple reaction monitoring (DMRM) with a cycle time of 500 ms and a retention time window of ±0.3 min. Two MRM transitions (quantifier and qualifier) were monitored for each compound—diazepam, nordazepam, oxazepam, and temazepam—to ensure selectivity and confirm identity; the specific precursor-to-product ion pairs, fragmentor voltages, and collision energies are listed in Table S1. Representative chromatograms illustrating the chromatographic resolution and retention times of all analytes in mouse spinal cord homogenate are shown in Figure S1. Data acquisition and processing were performed using Agilent MassHunter Quantitative Analysis software.

### 4.7. Statistical analysis

Only motoneurons with a resting membrane potential more negative than -50 mV and an overshooting action potential were kept for analysis. Only one cell was excluded from the dataset because its F-I gain was much higher than any other cell, and its identification as an alpha motoneuron was therefore questionable.

Note that it was not possible to blind the experimenter performing the experiments themselves, as the diazepam-treated animals required significantly higher amounts of anesthetics during the surgery and were therefore easily identifiable. However, the genotype and treatment information was obfuscated from the data files before data analysis and analyses were performed blind until all files were analyzed.

To assess the effects of genotype and chronic treatment on intrinsic motoneuron properties, each electrophysiological measure was analyzed using a linear mixed effects model with two fixed effects: genotype (wild type vs. mSOD1) and treatment (vehicle vs. diazepam), as well as their interaction (genotype × treatment). Because several motoneurons were recorded in each animal, and observations might not be strictly independent, mice were included as a random intercept. The model used was therefore:

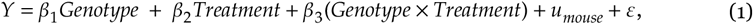

where Y is the property measured, *u*_*mouse*_ represents the random effect associated with each mouse and ε is the residual error.

To control for inflation of false positives arising from testing multiple properties, p-values were corrected for multiple comparisons using the Benjamini-Hochberg false discovery rate (FDR) procedure, with a significance threshold of q < 0.05. Given the a priori hypothesis that mSOD1 motoneurons would exhibit an exaggerated response to perturbation, genotype × treatment interaction effects were treated as the primary outcomes of interest, whereas main effects were considered secondary. Statistical significance was defined as an q-value (FDR-corrected p-value) less than 0.05.

All analyses were performed using Python v3.12, NumPy v.2.2.6 [49], Pandas v.2.2.3 [50], Seaborn v.0.13.2 [51], matplotlib v.3.10.7 [52], and statsmodels v.0.14.5 [53].

Proportions of synaptic inputs were analyzed using a multinomial mixed-effects model using the brms package [54] R v.4.6.0. Responses were categorized into discrete synaptic input types, and the probability of each category was modeled using a categorical (multinomial) likelihood with a logit link. Fixed effects included genotype, treatment, and their interaction (genotype × treatment). To account for repeated measurements and inter-individual variability, a random intercept for each mouse was included. Parameter estimates are reported as posterior means with 95% credible intervals. Evidence for effects was determined based on whether credible intervals excluded zero.

## Supplementary Materials

The following supporting information can be downloaded at: https://www.mdpi.com/article/doi/s1, Figure S1: Chromatographic retention times and MRM transitions for diazepam and its active metabolites in mouse spinal cord; Table S1: MRM parameters for diazepam and its metabolites.

## Author Contributions

Conceptualization, E.R. and M.M.; methodology, E.R., Y.C., D.L., and M.M.; software, M.M.; validation, E.R., Y.C., R.I., D.L., and M.M; formal analysis, M.M.; investigation, E.R., Y.C., R.I., and M.M.; data curation, M.M.; writing—original draft preparation, E.R., Y.C., and M.M.; writing— review and editing, E.R., Y.C., R.I., D.L., and M.M; visualization, E.R. and M.M.; supervision, D.L. and M.M.; project administration, M.M..; funding acquisition, M.M.

## Funding

This research was funded by NIH NINDS R01 NS110953.

## Institutional Review Board Statement

The animal study protocol was approved by the Institutional Ethics Committee of the University of Rhode Island (protocol AN2021-018).

## Data Availability Statement

The original data presented in the study are openly available in Zenodo at [DOI/URL].

## Acknowledgments

This research was supported in part using equipment and services available through the Rhode Island Institutional Development Award (IDeA) Network of Biomedical Research Excellence from the National Institute of General Medical Sciences of the National Institutes of Health under grant number P20GM103430.

## Conflicts of Interest

The authors declare no conflicts of interest. The funders had no role in the design of the study; in the collection, analyses, or interpretation of data; in the writing of the manuscript; or in the decision to publish the results.

## Abbreviations

The following abbreviations are used in this manuscript:

Adrb2: Beta-2-adrenergic receptor
Adrb3: Beta-3-adrenergic receptor
AHP: After-hyperpolarization
ALS: Amyotrophic lateral sclerosis
BH: Benjamini-Hochberg
DP: Deep peroneal
EDL: Extensor digitorum longus F frequency
FDR: False discovery rate
GABA: Gamma-aminobutyric acid I current
I_h_: Hyperpolarization-activated cation current
ΔI: Ioff-Ion
LC-MS/MS: Liquid chromatography with tandem mass spectrometry
PIC: Persistent inward current
PSP: Postsynaptic potential
mSOD1: Superoxide dismutase 1 (SOD1)^G93A^
MRM: Multiple reaction monitoring
mTOR: Target of rapamycin
SID: Once daily
SQ: Subcutaneous
T: Threshold
TA: Tibialis anterior
τ_m_: Membrane time constant
TS: Triceps surae
V: Voltage
WT: Wild type
IaIN: Ia inhibitory interneuron

## Notes

### Competing Interest Statement

The authors have declared no competing interest.

