## Supplemental Material for "Chronic diazepam reveals excessive homeostatic gain in SOD1^G93A^ mouse spinal motoneurons"

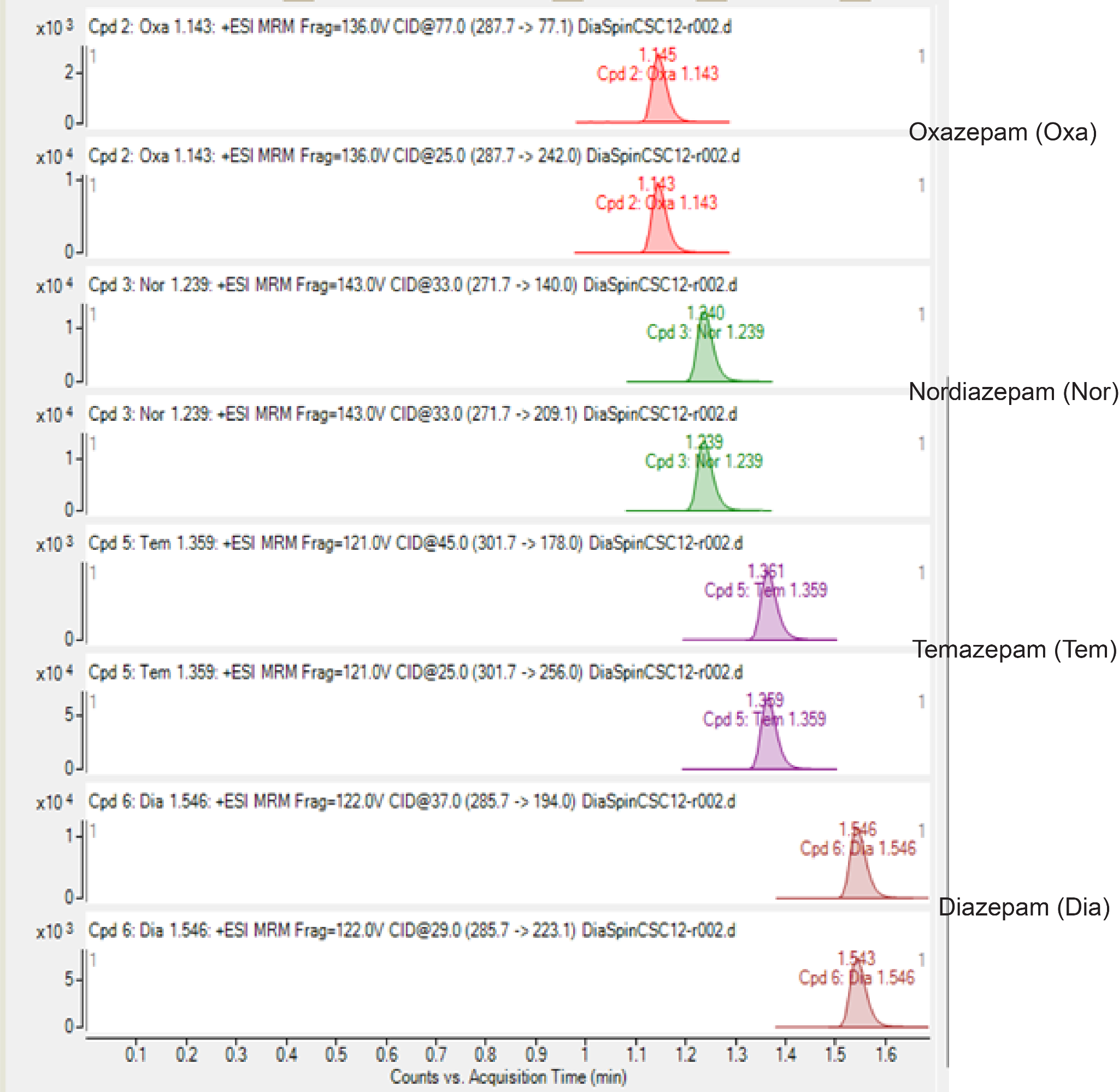

**Figure S1: Representative chromatograms and multiple reaction monitoring (MRM) transitions of diazepam and its active metabolites.** Chromatographic separation and detection were performed using a LC–MS/MS method operating in dynamic multiple reaction monitoring (DMRM) mode. Diazepam, nordazepam, oxazepam, and temazepam were monitored using two MRM transitions (quantifier and qualifier) to ensure selectivity and confirm analyte identity. The figure illustrates retention times and signal resolution of each analyte.

| **Compound** | **Precursor Ion (m/z)** | **Quantifier (m/z)** | **Qualifier (m/z)** | **Fragmentor Voltage (V)** | **Collision Energy (eV)** | **LLOQ (lower limit of quantification)** |
| --- | --- | --- | --- | --- | --- | --- |
| **Diazepam** | 285.7 | 223.1 | 194.0 | 122 | 29 / 37 | 0.36ng/ml |
| **Nordazepam** | 271.7 | 209.1 | 140.0 | 143 | 33 / 33 | 1.35ng/ml |
| **Oxazepam** | 287.7 | 242.0 | 77.1 | 136 | 25 / 77 | 0.36ng/ml |
| **Temazepam** | 301.7 | 256.0 | 178.0 | 121 | 25 / 45 | 1.50ng/ml |

**Table S1. Multiple reaction monitoring (MRM) parameters and lower limits of quantification (LLOQ) for diazepam and its metabolites.** The table summarizes the optimized MRM transitions, including precursor ions, product ions (quantifier and qualifier), fragmentor voltages, collision energies, and lower limits of quantification (LLOQ) for each analyte. The first MRM transition listed for each compound was used for quantification, while the second transition was used for confirmation of analyte identity. LLOQ values represent the lowest concentrations that could be reliably quantified with acceptable accuracy and precision under the validated LC–MS/MS conditions.
